# Morpho-functional analysis of the head glands in three Auchenorrhyncha species and their possible biological significance

**DOI:** 10.1101/2022.03.03.482260

**Authors:** Milos Sevarika, Andrea Di Giulio, Gabriele Rondoni, Eric Conti, Roberto Romani

**Author notes:** Corresponding author: Roberto Romani.

## Abstract

The Cicadomorpha *Philaenus spumarius, Neophilaenus campestris* and *Cicadella viridis* are known transmitters of the bacterium *Xylella fastidiosa*. Here, we studied the ultrastructural organization of their cephalic glands. Our investigations with scanning, transmission, FIB-SEM electron microscope and light microscope revealed for the first time in Auchenorrhyncha the presence of two types of cephalic glands. Both belonged to the class III epidermal glands, according to Noirot and Quennedey classification. Type I glands were the most common, being mainly located around antennae, lorum and gena. Moreover, these glands were observed also on the abdomen and thorax, always in association with sensilla trichoidea. The second type of glands were located exclusively at the apical part of the postclypeus in *P. spumarius* and *N. campestris.* The ultrastructural organization was similar in both types, being composed of a secretory cell and a conducting canal. Differences were observed in the width of the cuticular opening, being smaller in the type II glands. In addition, we have recorded the presence of a maxillary sensory pit in all species and described sensilla trichoidea ultrastructural organization. Finally, we discussed the ultrastructural organization of the glands and their potential biological role.

## Introduction

Glands are ubiquitous organs, found in most insect orders. They are known to have many different functions, including production of waxy-adhesive compounds used for egg attachment, attaching to the surface, building a cocoon, or restraining mating partners during copulation. Also, glands produce cleaning, defensive or pheromonal compounds, and have a role in protection against environmental factors or microorganisms (Betz, 2010; Di Giulio et al., 2009; Riolo et al., 2014). To date, exocrine glands have been recorded on antennae (Romani et al., 2005; Sevarika et al., 2021), head (Barbier et al., 1992), thorax (Billen & Ito, 2018), abdomen (De Santis et al., 2008) and legs (Zhang et al., 2021; Billen, 2009), showing clear functional localization among different species. Their relative number and abundance vary among species reaching the highest diversity in social insects that show a large repertoire of glands. For instance, in ants more than 40 anatomically distinct exocrine glands have been described until now (Hölldobler & Wilson, 2009; Billen, 2009).

Even though glands show great diversity as regards their position and cuticular release site organization, their ultrastructure is relatively uniform. Quennedey and Noirot classified glands into three groups based on their ultrastructural organization and on how the secretion crosses the cuticular barrier (Noirot & Quennedey, 1974). Glands of class I and III are commonly found among insects, while class II are less frequent and considered to be modified epidermal oenocytes (Noirot & Quennedey, 1974, 1991; Quennedey, 1998).

The presence of epidermal glands is a common feature in Hemiptera, being reported for several families either belonging to Heteroptera (true bugs) (Roell et al., 2020) or Homoptera (psyllids, aphids, scales) (Chen & Qiao, 2012; Ammar et al., 2015, 2013). For these insects, the role of the secretion is very diverse, including sexual communication (aphids and scales) (Boullis & Verheggen, 2016; Tabata & Ichiki, 2016), repellence or self-defence (true bugs) (Lockwood & Story, 1987; Krall et al., 1999), as well as release of wax or other protective material (scales, mealybugs) (Foldi & Pearce, 1985; Ahmad et al., 2020).

In the Auchenorrhyncha section, that includes cicadas, planthoppers and leafhoppers, besides wax glands the occurrence of exocrine glands is reported only in a few cases. For example, in *Xestocephalus subtessellatus* Linnavuori, (Cicadellidae), Cwikla and Freytag (1983) reported numerous pores present around eyes and antennae, with a potential secretory function. Similarly, on the anteclypeus and postclypeus of *Callitettix versicolor* (Fabricius) (Cercopidae), Liang (2020) reported two types of glands on males and one on females. Due to sex dimorphism, the author proposed that this latter type of gland in malescould be involved in the sex pheromone production used in intraspecific communication.

So far, all the studies were conducted solely by scanning electron microscope (SEM), thus they were unable to confirm the possible function of the investigated glands, nor to provide their ultrastructural organization. Here, we used the most common species of Aphrophoridae and Cicadellidae families as our model organisms. *Philaenus spumarius* L. and *Neophilaenus campestris* Fallén (Aphrophoridae) are spittlebugs known for the foam mass produced during preimaginal development, while *Cicadella viridis* L. (Cicadellidae) is characterized by its brochosomes (buckyball-shaped submicron proteinaceous particles, used as a superhydrophobic cover) production (Rakitov & Gorb, 2013; Rakitov, 2002). Moreover, two of these species (*P. spumarius* and *N. campestris*) are known as vectors of the bacterium *Xylella fastidiosa* Wells et al., whereas *C. viridis* is considered a potential vector (Janse & Obradovic, 2010; Cornara et al., 2019). Combining light, scanning (SEM) and transmission (TEM) electron microscope as well as focused-ion beam scanning electron microscope (FIB-SEM) we revealed for the first time detailed ultrastructural organization of the cephalic glands found in these species.

## Materials and methods

### Material examined

Adults of *P. spumarius, N. campestris* and *C. viridis* were collected by sweep-net samplings in meadows near Perugia (Central Italy). Once captured, insects were transferred in mesh cages (Kweekkooi 40×40×60 cm, Vermandel, Hulst, The Netherlands) and maintained under controlled environmental conditions (25±2 °C, L16:D8, RH 75±5%). *Philaenus spumarius* and *C. viridis* were reared and maintained on 1- to 3-week-old potted plants of *Vicia faba minor* L., whereas *N. campestris* was provided with 20-cm-tall *Triticum aestivum* L. plants grown on small trays (10×20 cm).

### Scanning Electron Microscope (SEM)

Five individuals of each sex for each species were observed under scanning electron microscope. Prior to their preparation, individuals were anesthetized by low-temperature exposure (−18°C for 2 min) and placed in 50% ethanol. Under a stereomicroscope, the heads were detached from the thorax by micro scissors. In the case of *C. viridis*, the heads were cleaned from the wax and brochosomes through over-night submersion into 5% KOH, conducted at room temperature. Next, heads were washed with bidistilled water and treated with pure glacial acetic acid for 5 min, and then washed by bidistilled water three times for 5 min. Subsequently, heads were dehydrated in a series of graded ethanol (60, 70, 80, 90, 95, 99%, each step for 15 min, except the last which was repeated twice), then exposed to Hexamethyldisilazane (HDMS, Sigma-Aldrich, Dorset, UK) and allowed to dry under a ventilated hood at room temperature. Finally, the samples were mounted on aluminum stubs, and gold-coated using a “Balzers Union^®^ SCD 040” unit (Balzers, Vaduz, Liechtenstein).

To observe glands from the internal part of the head, prior to desiccation, the head was cut in half longitudinally by micro scissors end submerged into a 5% KOH/water solution at 50°C for 10 min. This enabled to clean the internal soft tissues and clearly visualize gland position and internal cuticular surface. Later, specimens were dehydrated and gold-coated as described above. The observations were carried out using a FE-SEM Zeiss^®^ SUPRA 40 (Carl Zeiss NTS GmbH, Oberkochen, Germany) and a Philips^®^ XL 30 (Eindhoven, The Netherlands) operating at 7-10 KV, WD 9-10 mm and analyzed by a SMART-SEM^®^ software.

### Light and Transmission Electron Microscope (TEM)

Five individuals of each sex and species were anesthetized by exposure to low-temperature (−18°C for 2 min), and soon after they were immersed in a solution of 2% glutaraldehyde and 2.5% paraformaldehyde in 0.1 M cacodylate buffer+5% sucrose, at pH 7.2–7.3. Then, heads were detached from the base, cut into small pieces to facilitate penetration of the fixative, and left at 4°C overnight. After rinsing in the buffer (2×15 min), the specimens were post-fixed in 1% OsO4 (osmium tetroxide) for 1 h at 4 °C, then rinsed in the same buffer (2×15 min). The specimens were dehydrated in an ascending series of graded ethanol from 60 to 99% and embedded in Epon-Araldite resin with propylene oxide as bridging solvent. Semi-thin sections (0.5 µm) were taken with glass-knife on a 2188 Ultratome Nova ultramicrotome (LKB^®,^ Stockholm, Sweden) and stained with toluidine blue. The observations were conducted on Zeiss light microscope (Zeiss 47 30 12-9902, Germany) with the Ph2 Neofluar 16x objective (Zeiss, Germany). For the transmission electron microscope, the sections were taken with the diamond knife and mounted on formvar-coated 50 mesh grids. The sections on grids were stained with uranyl acetate (20 min, room temperature) and lead citrate (5 min, room temperature). Finally, the sections were observed with a Philips^®^ EM 208. Digital pictures (1376 × 1032 pixels, 8b, uncompressed greyscale TIFF files) were obtained using a high-resolution digital camera MegaViewIII (SIS^®^) connected to the TEM.

### Focused Ion Beam – Scanning Electron Microscope (FIB-SEM)

Five individuals of each sex of *C. viridis* were processed, as described above for the other species, up to the post-fixation step with OsO4. After post-fixation, specimens were en-bloc stained with 2% aqueous uranyl acetate. Subsequently, the samples were dehydrated in a graded ethanol series (70%, 85%, 95%, 30 min each and 100% for 2 h), embedded in Epon-Araldite resin and finally polymerized for 72 h at 60 °C. In order to acquire ultrastructural information, the resin-embedded samples were cut into sequential slices, 15–20 μm thick, using the aforementioned ultramicrotome equipped with a glass knife. Thick slices were attached to aluminum stubs with a conductive adhesive carbon disk, sputtered with a thin layer (30 nm) of gold in a K550 sputter coater (Emithech, Kent, UK), and analyzed with FIB/SEM following the ‘Slice & Mill’ method (Di Giulio & Muzzi, 2018). This allows milling of regions of interest, by using an ion beam, and imaging of ultrastructural details present on the freshly milled surface, by detecting secondary electrons.

## Results

The general structure of the head was similar among the investigated species, with a typical sub-triangular shape (when observed looking directly at the frons) (Figure 1A, 2A, 7A). The top of the heads appeared quite smooth and devoid of cuticular structures (Figures 2B, C). In *P. spumarius* and *N. campestris,* the head was clearly divided into different sclerites by pronounced sutures. A difference between the two species was observed regards the position of compound eyes. These were well developed in both species, but in *P. spumarius* they were positioned more anteriorly with respect to *N. campestris* (considering the head in natural position) (Figures 1A, 2A). In both species the head size was similar; in *P. spumarius* was about 1200 µm widt and 1600 µm length, while in *N. campestris* width was 1100 µm and length 1750 µm). The anteclypeus and postclypeus encompassed the central portion of the head. On postclypeus, a double series of furrows was observed, with the space between them covered by numerous sensilla trichoidea (Figures 1C, 1F). Sensilla trichoidea were abundant on all major sclerites: anteclypeus, postclypeus, lorum, and gena. A series of rounded cuticular openings (about 2.5 µm in diameter) were found around sensilla trichoidea (Figures 1C, 2F, 6A). These openings were denoted as gland type I.

**Figure 1.**
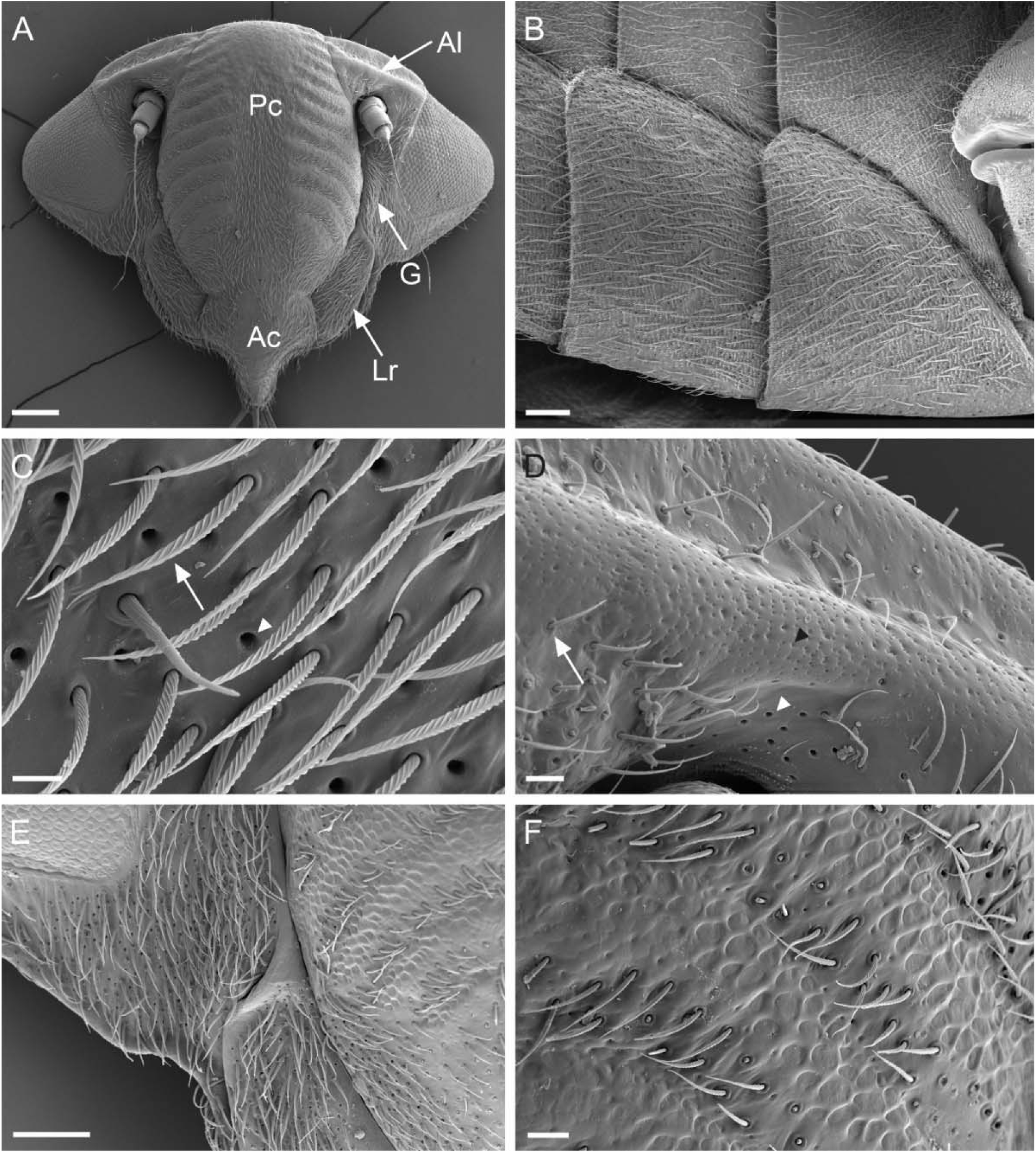
Scanning electron micrographs of *Philaenus spumarius* female A) General view of the head. B) Abdominal sternites showing numerous glands type I around sensilla trichoidea. C) Distribution of the gland type I (arrowhead) around sensilla trichoidea (arrow). D) SEM micrographs showing a portion of the antennal ledge with the distribution of the glands type I (white arrowhead) and type II (black arrowhead). The arrows point to the sensilla trichoidea, which are less abundant on the antennal ledge. E) Distribution of the numerous glands type I on the lorum and gena in P. spumarius. F) The zone on a postclypeus with the sensilla trichoida distribution. AL antennal ledge, G gena, Lr lorum, Ac anteclypeus, PC postclypeus. Scale bars A = 200 µm, B =15 µm, C =10 µm, D =20 µm, E =100 µm, F =20 µm.

**Figure 2.**
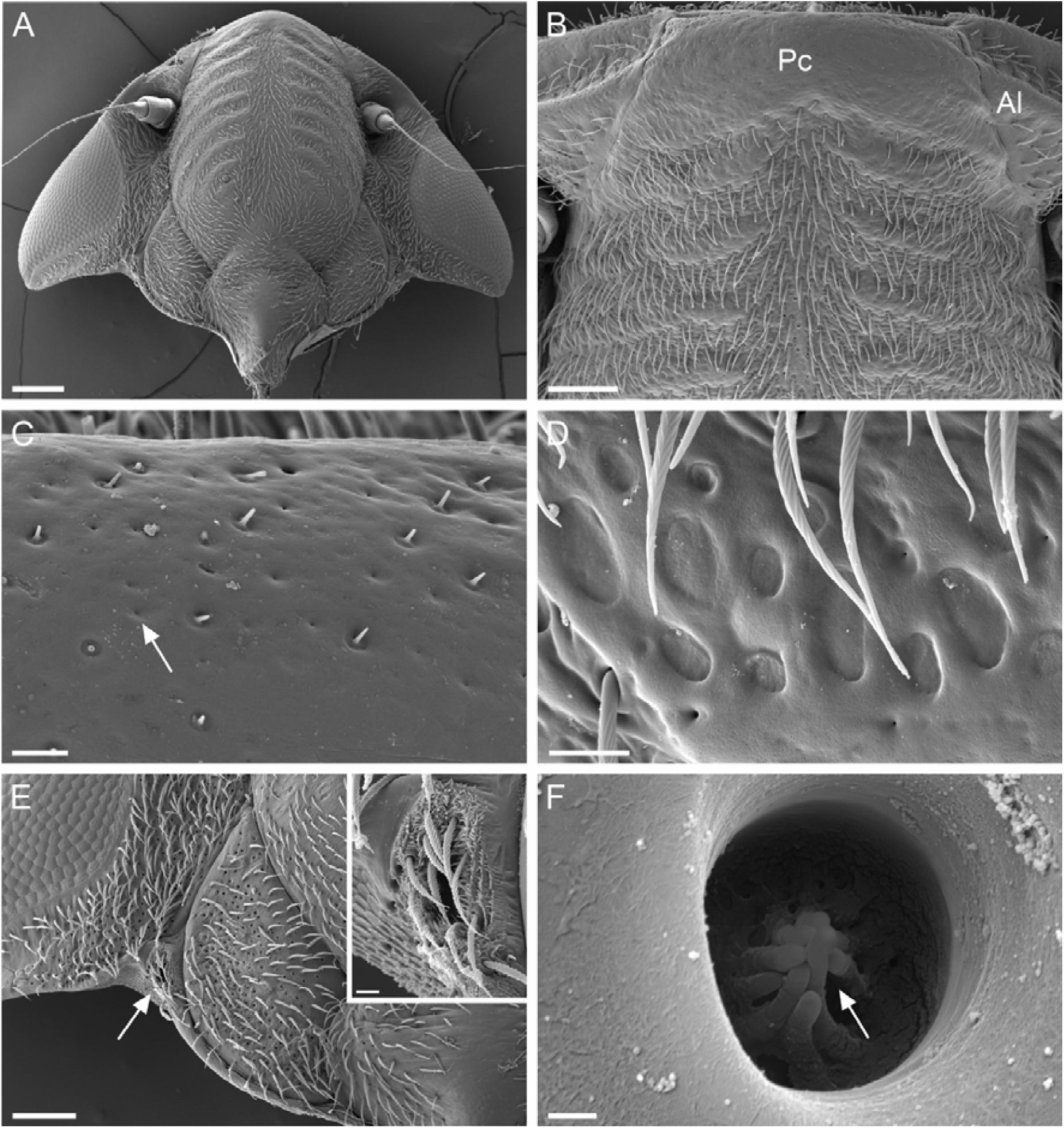
Scanning electron micrographs of *Neophilaenus campestris* female. A) General view of the head. B) Higher magnification of the postclypeus with difference in the sensilla trichoidea localization. Note, the apical part of the postclypeus is devoid of sensilla trichoidea. C) Small cuticular openings indicating presence of the glands type II (arrow). D) SEM micrograph showing the zone on postclypeus, between two lines of furrows with the sensilla trichoidea. E) Higher magnification on the region between lorum and gena. Maxillary sensory pit is covered by sensilla trichoidea (see inset). F) View on the gland type I chamber, showing finger-like projections (arrow). Ac anteclypeus, Pc postclypeus. Scale bar A = 200 µm, B = 100 µm, C, D = 10 µm, E = 100 µm, imprint = 10 µm, F = 300 nm.

From the antennal ledge, a decrease in the number of sensilla trichoidea was observed, with the apical part of the anteclypeus being devoid of them (Figures 1D, 6C). This area was characterized by the presence of smaller cuticular openings (about 0.3 µm in diameter), denoted as gland type II (Figures 2C). Moreover, on the maxillary plate, a sensory pit was recorded, which was surrounded by numerous sensilla trichoidea (Figures 1E, 2E). Within it, a small structure covered by numerous scales was observed (Figures 2F).

In *C. viridis* the head showed the same general features reported for the two spittlebug species (Figure 7A). However, the head did not show the presence of sensilla trichoidea, while abundant brochosomes were present, covering the cuticle (this could be observed in non-cleared specimens; Figures 7A, 7F). On the head, peculiar cuticular openings were found. These structures were denoted as gland type I, located particularly on the areas around the antenna and on the gena (Figures 7B-D). The maxillary sensory pit was located on the maxillary plate. The area surrounding the pit was devoid of sensilla and appeared rather as a hollow structure, without a specific sensory organization within it (Figure 7E).

### Gland type I

Gland type I was the most abundant on *P. spumarius* and *N. campestris.* They were found on head, abdomen, and thorax, always in association with sensilla trichoidea (Figures 1A-F, 2A-B, F). Their distribution was random on different segments. On the head, higher glands density was recorded on the lorum and gena. Externally, the glands were characterized by a rounded cuticular opening (in *P. spumarius* and *N. campestris* the diameter was about 2.5 µm, while for *C. viridis* it was about 7 µm) with finger-like projections. Such projections in *C. viridis* were in line with the cuticle, while, for *P. spumarius* and *N. campestris* were found at the base of the glandular pit (Figures 2F, 7D). Internal observation after KOH digestion revealed the presence of a short conducting canal (Figure 6B). The innermost part of the conducting canal presents the remnants of the end apparatus. TEM investigations revealed that these glands were composed of 2 cells: the secretory cells, which form the end apparatus and synthesize secretory products, and the duct cell that produces the evacuating duct (Figures 3A-C, 4A-D). The secretory cell contained the large nucleus and groups of electron-dense secretory vesicles. The end apparatus was rounded and highly microvillated, occupying the largest part of the cell. The secretory products of the cell accumulated at the base of the end apparatus and were secreted through the conducting canal (Figures 8C-E). The conducting canal had a constant diameter and consisted in a thin layer of epicuticle lining a thicker endocuticle (Figures 4A-B, 8D-G). In our observations, the ducts appeared to be tightly closed because of the position of the internal epicuticle. The length was higher in *P. spumarius* compared to *N. campestris* (Figures 3A, 4A). At the gland surface, secretory traces were observed (Figure 4A-B).

**Figure 3.**
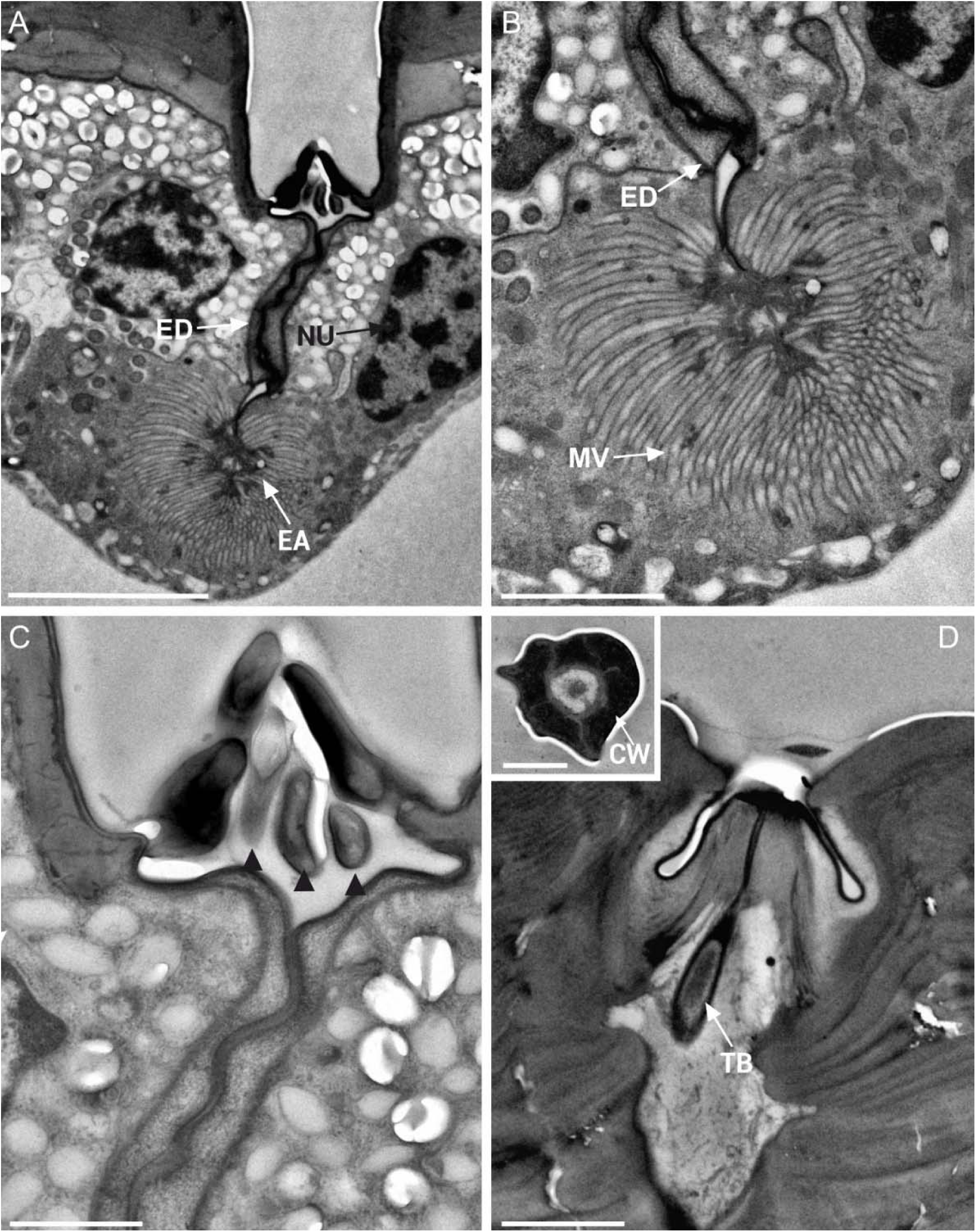
Transmission electron micrograph of gland type I and sensilla trichoidea in *Neophilaenus campestris* female. A) Longitudinal section of gland type I showing end-apparatus (EA) and evacuating duct (ED). On tip of the evacuating duct a finger-like ridges can be seen. NU nucleus. B) Higher magnification of the end-apparatus showing its highly microvillated structure (MV) and starting point of the evacuating duct (ED). C) Closer look at the finer-like projections inside the glandular pit, covering the exit point of the evacuating duct. D) Longitudinal TEM section through the sensilla trichoidea showing tubular body (TB). Inset shows thick cuticular wall (CW) of sensilla trichoidea. A = 5 µm, B = 2 µm, C = 10 µm, D = 2 µm, Inset = 1 µm.

**Figure 4.**
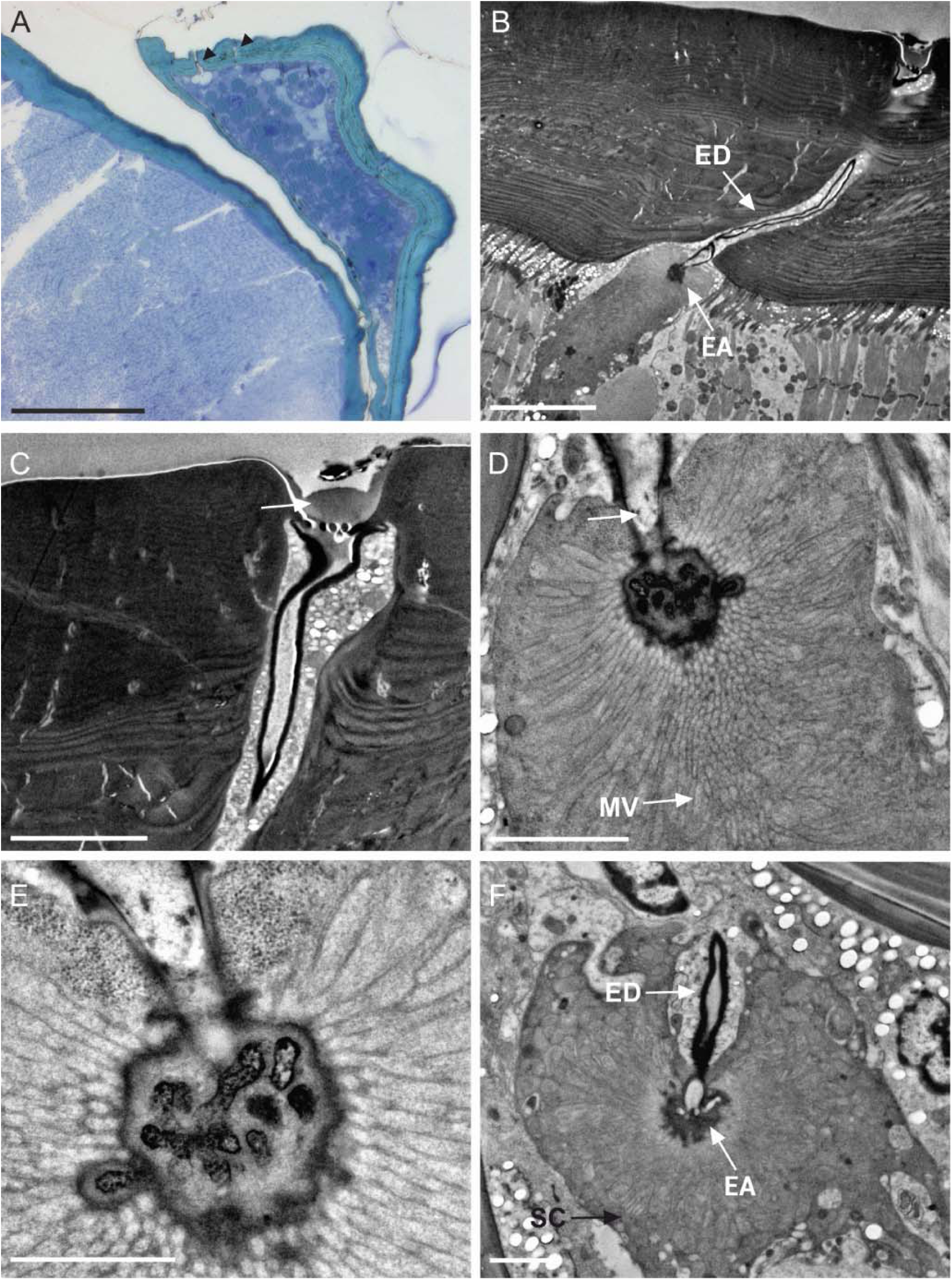
Light and transmission electron micrograph of gland type I in *Philaenus spumarius* female. A) Semi-thin section taken at the antennal-ledge level showing position of the cuticular openings (black arrowheads) B) Longitudinal TEM section showing end-apparatus and evacuating duct of the gland type I. C) Higher magnification at the glandular opening revealing the glandular products located at the base of the gland base. D) End-apparatus of the gland type I with the duct cell (arrow) and highly microvillated end-apparatus. E) closer look at the duct cell from D. F) Secretory cell of gland type I with the end-apparatus and evacuating duct. SC secretory cell, EA end-apparatus, EV evacuating duct, MV microvilli Scale bars: A = 100 µm, B = 10 µm, C = 5 µm, D = 2 µm, E = 1 µm, F = 2 µm.

**Figure 5.**
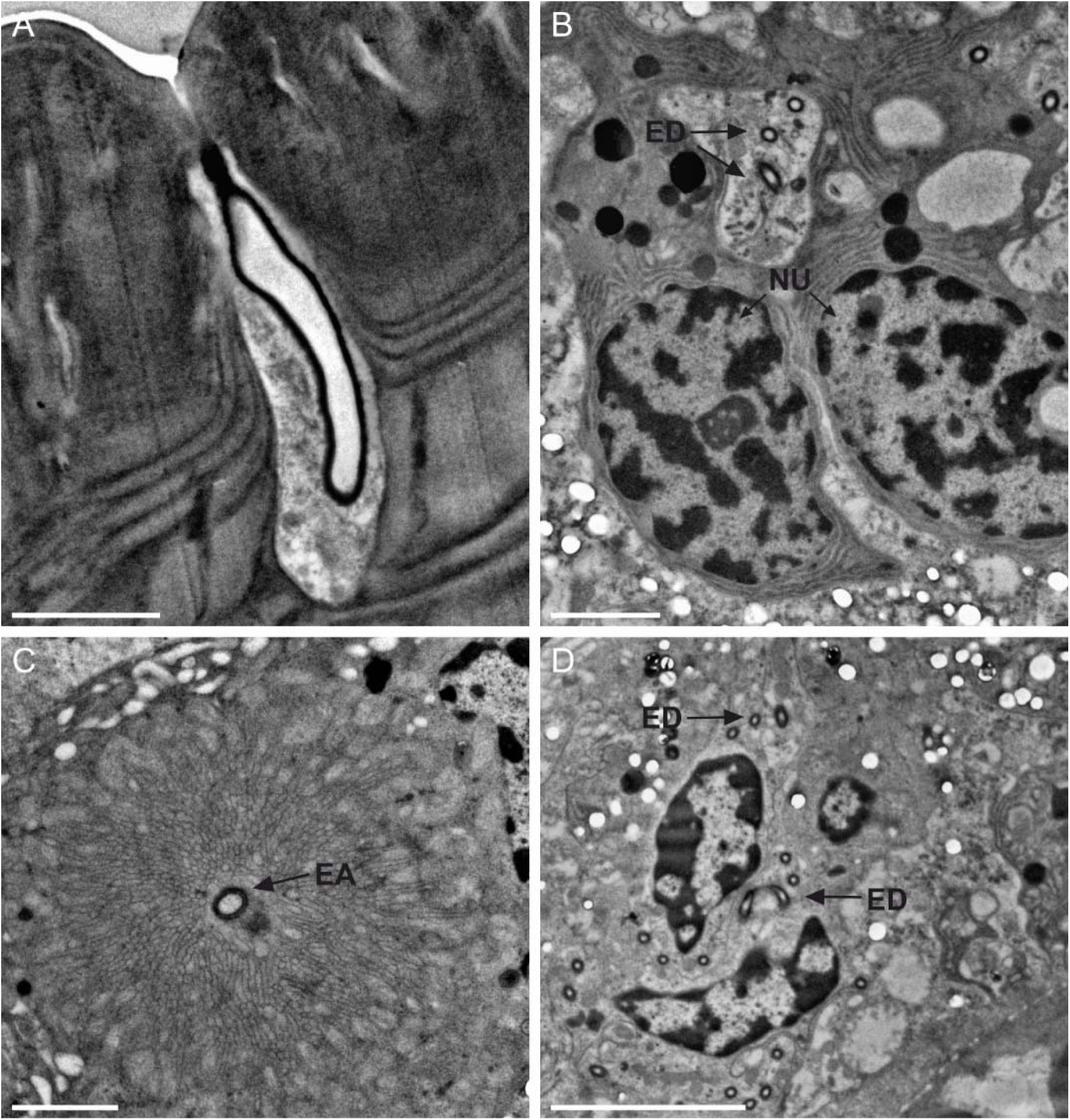
Transmission electron micrographs of the gland type II in *Philaenus spumarius* female. A) Longitudinal section through the apical part of the gland, showing the evacuating duct with a small glandular opening. B) Numerous segments of the evacuating duct around two large nuclei. C) Cross section of the secretory cell showing the evacuating duct surrounded by numerous microvilli belonging to the end-apparatus. D) Network of the evacuating duct around two large nuclei. ED evacuating duct, EA end-apparatus, Scale bars: A, B, C = 2 µm, D = 5 µm.

**Figure 6.**
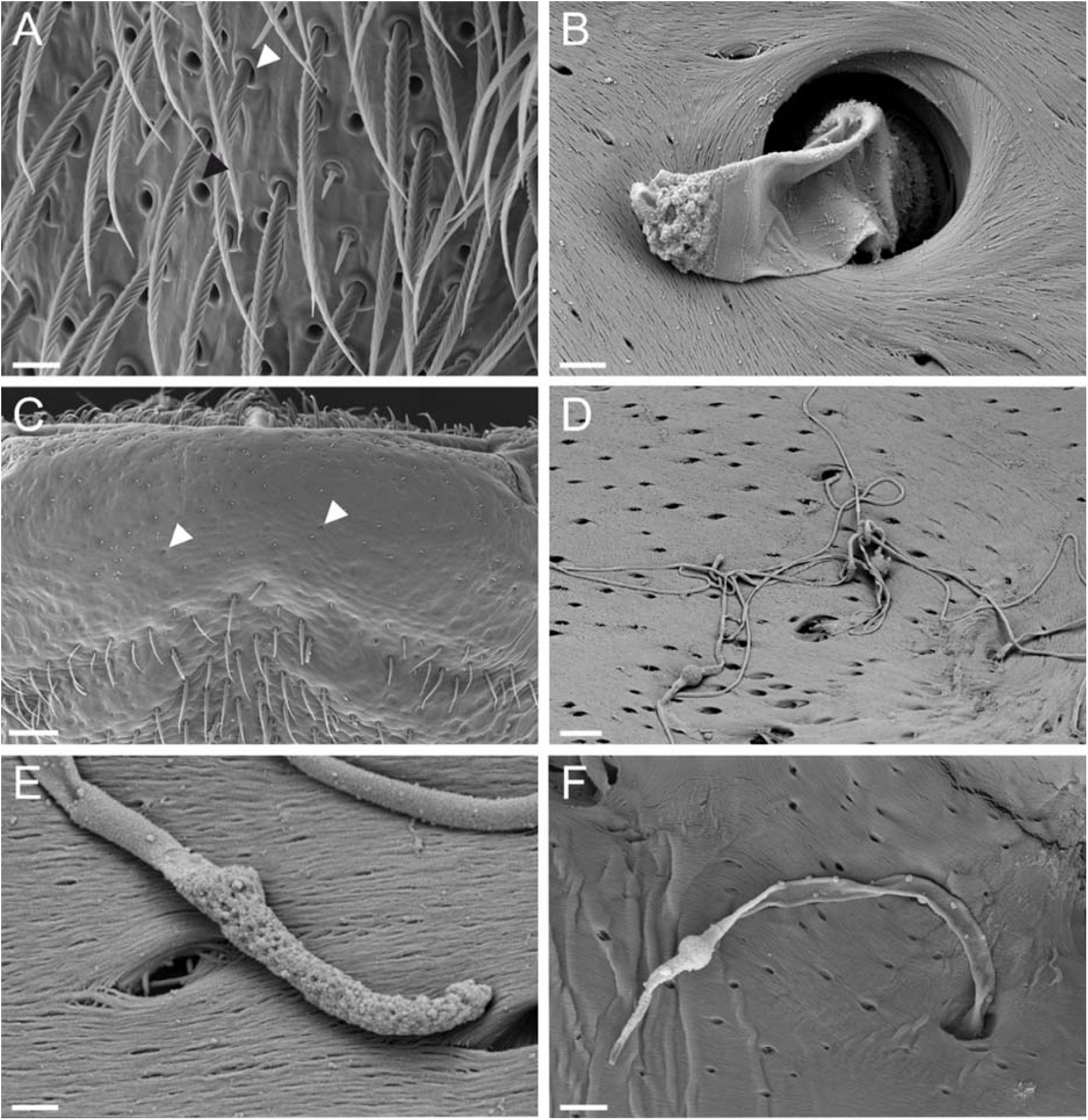
*Philaenus spumarius* female scanning electron micrographs of the glands type I and type II external and internal view. A) Numerous glands type I (black arrowhead) located around sensilla trichoidea (white arrowhead). B) Gland type I from the internal part of the head. This gland type is characterized with the short and wide evacuating duct. C) Distribution and cuticular openings of the glands type II. D, E, F) The gland type II from the internal part of the head showing long evacuating duct which are often interconnected. On E and F different shape of the evacuating duct can be observed. Scale bars: A = 10 µm, B = 1 µm, C = 40 µm, D = 3 µm, E = 0.4 µm, F = 2 µm.

**Figure 7.**
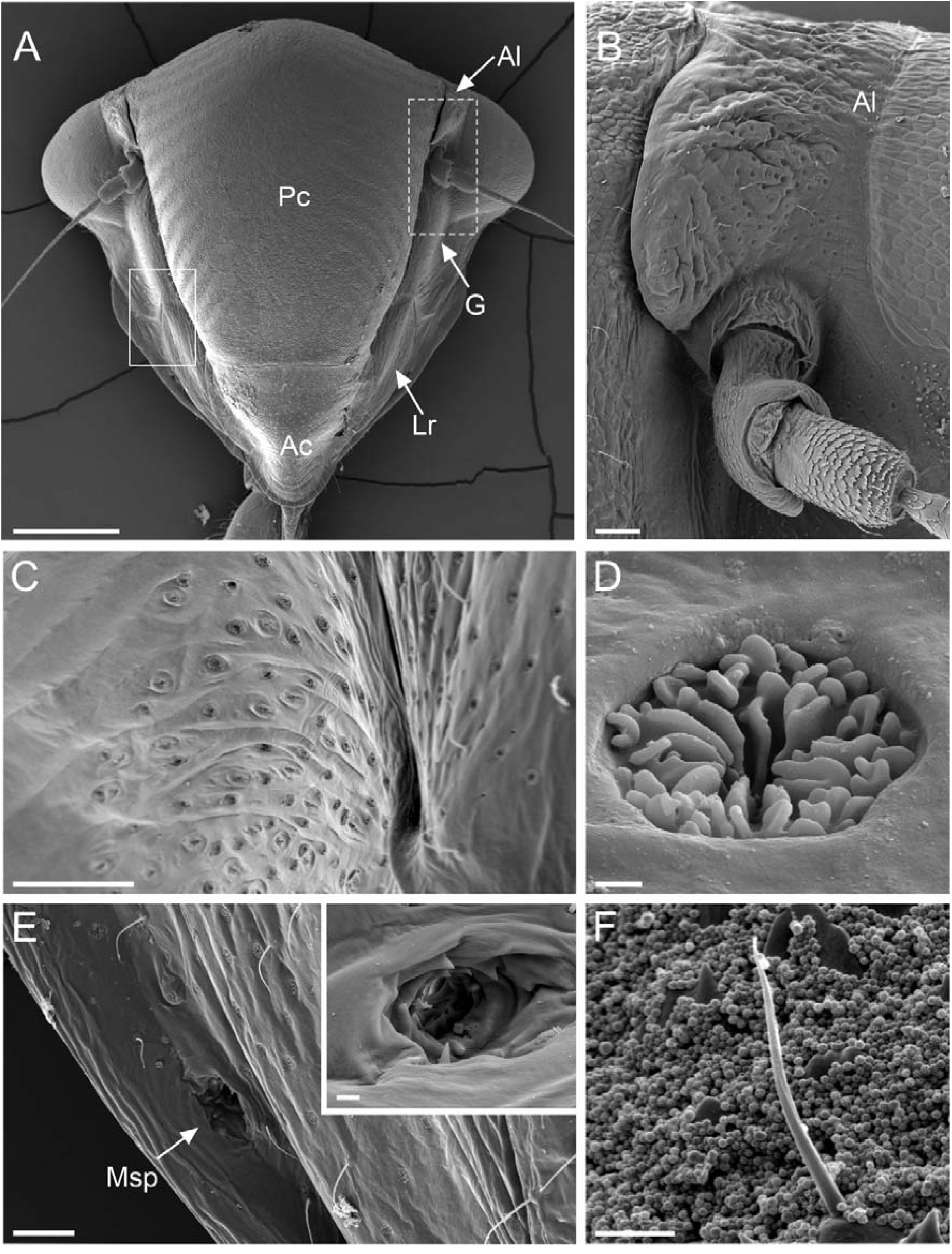
Scanning electron micrographs of *Cicadella viridis* female. A) General view on the head. Full-line rectangle marks transient zone between lorum and gena, while rectangle with broken lines indicate the zone around the antennae. B) Higher magnification from the picture A, showing numerous glands around the antennae and the antennal ledge. C) Magnification from the picture A, with the numerous cuticular pores. D) High magnification of the finger-like projections of the gland type I located at the cuticular base. E) Maxillary sensory pit (Msp) without sensilla trichoidea around it. Inset – Magnification of the maxillary sensory pit, showing its hollow structure. F) View on the C. viridis prior cleaning process, showing numerous brochosomes covering the cuticle. Scale bars: A, B = 40 µm, C = 50 µm, D = 1 µm, E = 30 µm, Inset = 4 µm, F = 5 µm.

**Figure 8.**
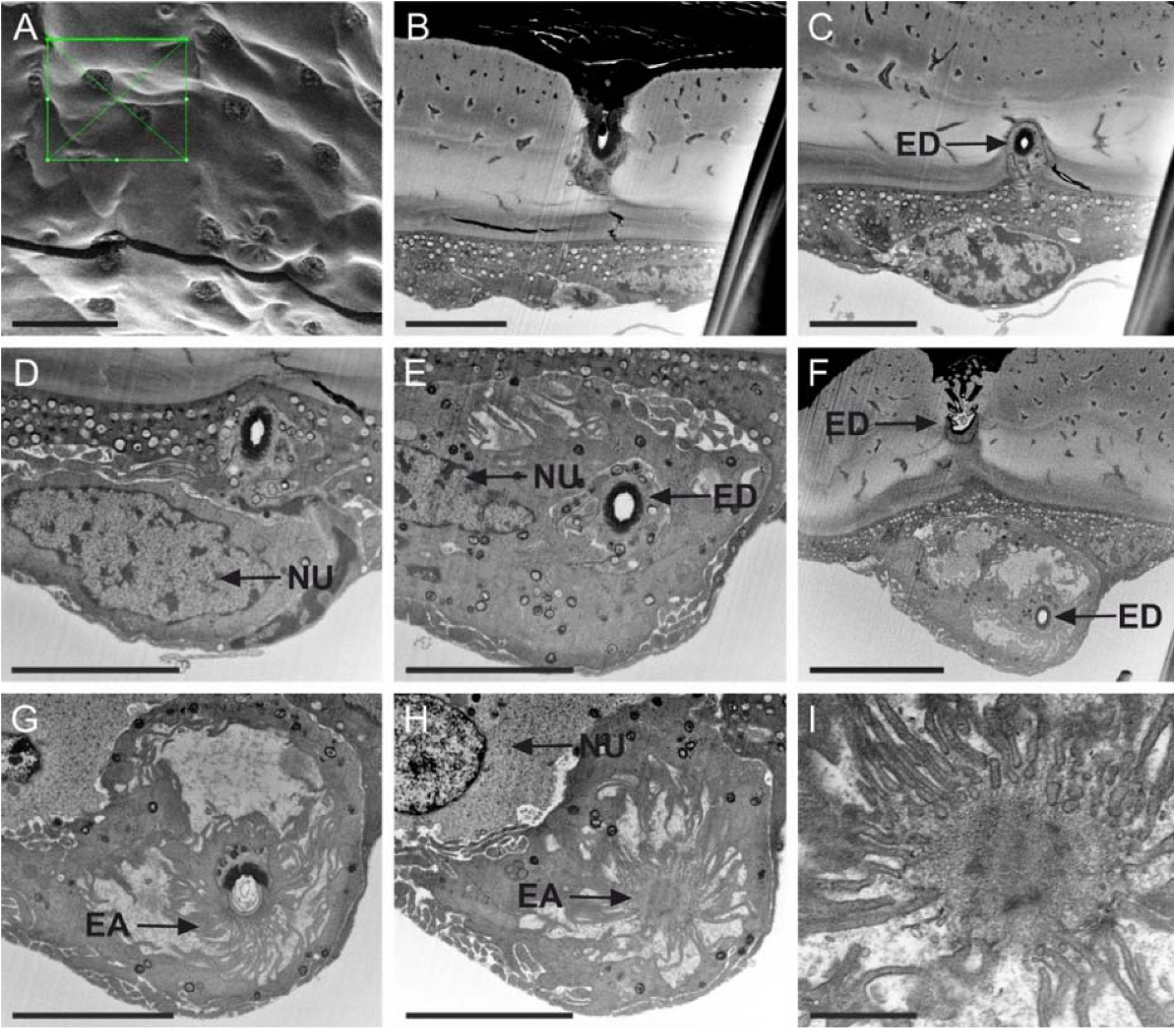
Images of *Cicadella viridis* female gland type I taken with focused ion-beam microscope. A-I) Ultrastructural organization of the gland type I taken at various depths. On the picture A, the green rectangle indicates the zone that was milled. Through the different pictures, large nucleus (NU) can be observed. Evacuating duct can be seen from the gland initial point (pictures B to E), while, from the picture G, an initial zone of the secretory cell can be seen. Scale bars: A = 20 µm, B, C, D, E, G, H = 5 µm, F = 10 µm, I = 1 µm.

### Gland type II

The second type of glands were observed solely at the apical part of the postclypeus and antennal ledge in *P. spumarius* and *N. campestris*, although we characterized ultrastructure only those of *P. campestris* (Figures 1A, 2B). The cuticular opening was small, about 0.3 µm in diameter. This opening is internally connected to a conducting canal, which was narrower and longer, compared to the duct observed in gland type I (Figures 6C-F). The ultrastructural organization revealed also in this case the presence of bicellular units composed of a secretory cell and a duct cell (Figures 5A-D). Due to their length, the conducting canals of different cells were interconnected creating a network of duct tracts. At the end of the duct, an end-apparatus of a different shape was detected (Figures 6D-F). This gland lacked the finger-like protrusions at the level of the external opening, and the conducting canal was made of epicuticle only.

### Sensilla trichoidea

Sensilla trichoidea were abundant on *P. spumarius* and *N. campestris* head. The sensilla had a distinct socket (Figures 1C). The external cuticular wall had peculiar ridges, which appeared in sequence, directed to the median line of the sensilla (Figures 1C). The sensillum was 45 µm long on average, and exhibited a pointed tip. The cross-section taken at the apical part of the sensilla showed a thick cuticular wall without sensory neurons inside. The TEM pictures taken at the basal part of the sensilla revealed the presence of a sensory neuron with a tubular body at its base (Figures 3D).

## Discussion

In our study we investigated the presence of epidermal glands located on the head of two spittlebugs and a leafhopper species, all belonging to the large systematic group of Auchenorrhyncha. In all species, we found external cuticular openings located on various head regions, mainly the anteclypeus, the areas around the eyes, the genae and the head vertex. These openings are internally connected to secretory units that, based on morphological features, correspond to Class III secretory cells, according to the classification proposed by Noirot and Quennedey (Noirot & Quennedey, 1974). The Class III glands are frequently described in insects. Usually, they are composed of one or more secretory cells associated with one or two duct cells producing the conducting canals (Bin et al., 1999; Bartlet et al., 1994; Barbier et al., 1992; Giglio et al., 2005; Di Giulio et al., 2009). In our model species, we described two types of exocrine glands. Both types of glands reported here were of the simple form, composed of one secretory cell and a conducting canal. They differed in their structural organisation and localisation. Glands of type I of all species were abundant on the same head sclerites. Externally, in *C. viridis* the finger-like projections were in the line with the cuticle, while, in *P. spumarius* and *N. campestris* they were found at the gland chamber base. Functionally speaking, the finger-like projections could protect glands from being filled with dirt, thus blocking compounds release from the conducting canal. Contrary to glands type I, glands type II were located solely at the apical part of the postclypeus. As a result of small cuticular openings, these glands did not have finger-like projections. Ultrastructurally, the conducting canal was significantly longer than in Glands type I, creating a network of conducting canals when looked from the internal part of the head.

Until now, in Auchenorrhyncha, several studies reported presence of structures for which a glandular function was hypothesized or, at least based on external morphology, it could be speculated. For instance, in *Xestocephalus subtessellatus* Linnavuori, (Cicadellidae), Cwikla and Freytag (1983) hypothesized a secretory function of a reported structure. In *Callitettix versicolor* (Fabricius) (Cercopidae), Liang (2020) reported the presence of putative exocrine glands located on male head, which, potentially could produce sex pheromones. Moreover, according to Rakitov (2009), the structure denoted as coeloconic sensillum could possibly be the external opening of an integumental gland. In addition, small cuticular openings in the area around antennae were observed in *Isobium gibbosum* Melichar, *Globularica diversicolorata* (Stroiński et al., 2011) *Cyamosa camelouca* (both Fulgoromorpha) (Stroiński et al., 2011).

The glands described here showed localization and ultrastructural difference, which could lead to difference in their function. Glands type II were found on frons, without any structures around them, thus proving possibility of a clear contact with other individuals. In the case of *C. versicolor*, specific behaviour during the courtship is exhibited, during which, the male positions his leg on the female’s frons (Liang, 2020). Thus, we hypothesize that these glands could produce compounds used in intraspecific communication. Glands type I were the most abundant glands, being found on the head as well as on the abdomen. For these glands, we exclude wax production, since these species are not wax-producing insects, conversely to other species belonging to Fulgoroidea, such as *Metcalfa pruinosa* (Say) (Lucchi & Mazzoni, 2004). Based on the traces we found at the glandular base, indicating the low volatility of the released compounds, and their widespread distribution through the body, we hypothesize these glands could produce compounds utilized in different possible biological contexts (cleaning substances, intraspecific communication etc). The release of these compounds on the substrate is probably facilitated by the abdomen trembling produced during vibrational communication (Laumann et al., 2013). Nevertheless, the exact function of the produced compounds needs to be confirmed by further behavioural and chemical analysis.

Apart from glands, we observed Evan’s organ (also called maxillary sensory pit) on the maxillary plate. Evan’s organ is common in Auchenorrhyncha and Coleorrhyncha and absent in Sternorrhyncha and Hemiptera (Evans, 1973; Wang et al., 2013). Indeed, until now it has been reported in *Zema gressitti* Fennah (Fulgoromorpha) (Wang et al., 2013), *Borysthenes* (Fulgoromorpha) (Liang, 2005), *Cixiopsis punctatus* Matsumura (Fulgoromorpha) (Wang et al., 2014), several Cicadelloidea (Evans, 1975), *Zyginidia pullula* (Boheman) (Typhlocybinae), *Empoasca vitis* (Goethe) (Typhlocybinae) and *Graphocephala fennabi* Young (Cicadellinae) (Tavella & Arzone, 1993). Evan’s organ is reported as a sensory pit containing a palp-like process or finger-like lobe. Based on its multiporous structure, an olfactory sensory function was proposed (Wang et al., 2013). In our study, we did not perform an in-depth analysis of the maxillary sensory pit to shed light on its real structural organization, which could be related to sensorial but also secretory function. This will be the subject of a coming study.

*Philaenus spumarius* and *N. campestris* heads were abundantly covered with sensilla trichoidea. The cuticular wall was of peculiar shape, not commonly found in other species. The ultrastructural organization was general, being structured of a tubular body located at the base and having aporous cuticular wall (Sevarika et al., 2021; Riolo et al., 2016). As a result of its ultrastructural organization, a mechanoreceptive role was given to this sensillum.

Here we presented the first detailed ultrastructural organization of cephalic glands in Cicadomorpha species. Both glands belonged to the class III by Noirot and Quennedey classification, which are often associated with pheromone production. Recently, in *Philaenus spumarius*, and for the first time in Cercopoidea, presence of a potential sex-pheromone was reported (Sevarika et al., under review). Thus, this study contributes to filling the gap about the involvement of chemical compounds in intraspecific communication in this group of insects. Furthermore, the future study should aim to elucidate the chemical composition of gland products, as well as how individual components affect behavioural ecology in these insects. Moreover, it would be of interest to understand how widespread is gland presence among other Auchenorrhyncha.

## Acknowledgements

Scanning electron microscopy, transmission electron microscopy and FIB data were obtained at the Centro Universitario di Microscopia Elettronica e a Fluorescenza (CUMEF; University of Perugia) (SEM and TEM) and Laboratorio Interdipartimentale di Microscopia Elettronica (LIME, University Roma Tre) (SEM and FIB).

